# FITdb, an Integrated Functional Immunogenomics and Transcriptomics Database

**DOI:** 10.64898/2026.07.28.741304

**Authors:** Xinjian Cen, Qixuan Ma, Kevin Kim, Susanne Gamas-Vis, Ananda Goldrath, Maximilian Heeg, Miguel Reina-Campos

## Abstract

Genetic screens in immune cells enable the systematic interrogation of gene function at scale, uncovering key regulators of cell functions such as tumor cell killing and persistence. However, existing datasets typically focus on specific biological questions, employ targeted gene panels, are generated under diverse experimental conditions, and are not readily accessible, which together limit their integration and future usability. To address this, we developed the Functional Immunogenomics and Transcriptomics Database (FITdb), a freely accessible resource that harmonizes functional genomics datasets for the study of immune cell biology. FITdb currently integrates 43 independent functional genetics screens, including 32 pooled and 11 single-cell screens, spanning 20, 696 mouse genes and 22, 293 human genes across 199 immune cell types and conditions. All datasets are uniformly re-analyzed to enable cross-study comparisons. FITdb provides intuitive, gene-centric visualizations, detailed exploration of individual screens, and access to sgRNA-level data. Additionally, built-in tools such as “*Compare MyGeneSet*” and “*Compare MyScreen*” identify statistically significant overlaps between user-defined gene lists and functional gene sets in FITdb, and enable direct comparison of user-generated screening data with existing datasets, respectively. Together, FITdb provides a comprehensive, user-friendly platform for accelerating the discovery of immune regulatory programs. The database is freely available at https://fitdb.lji.org.

**Graphical Abstract:** 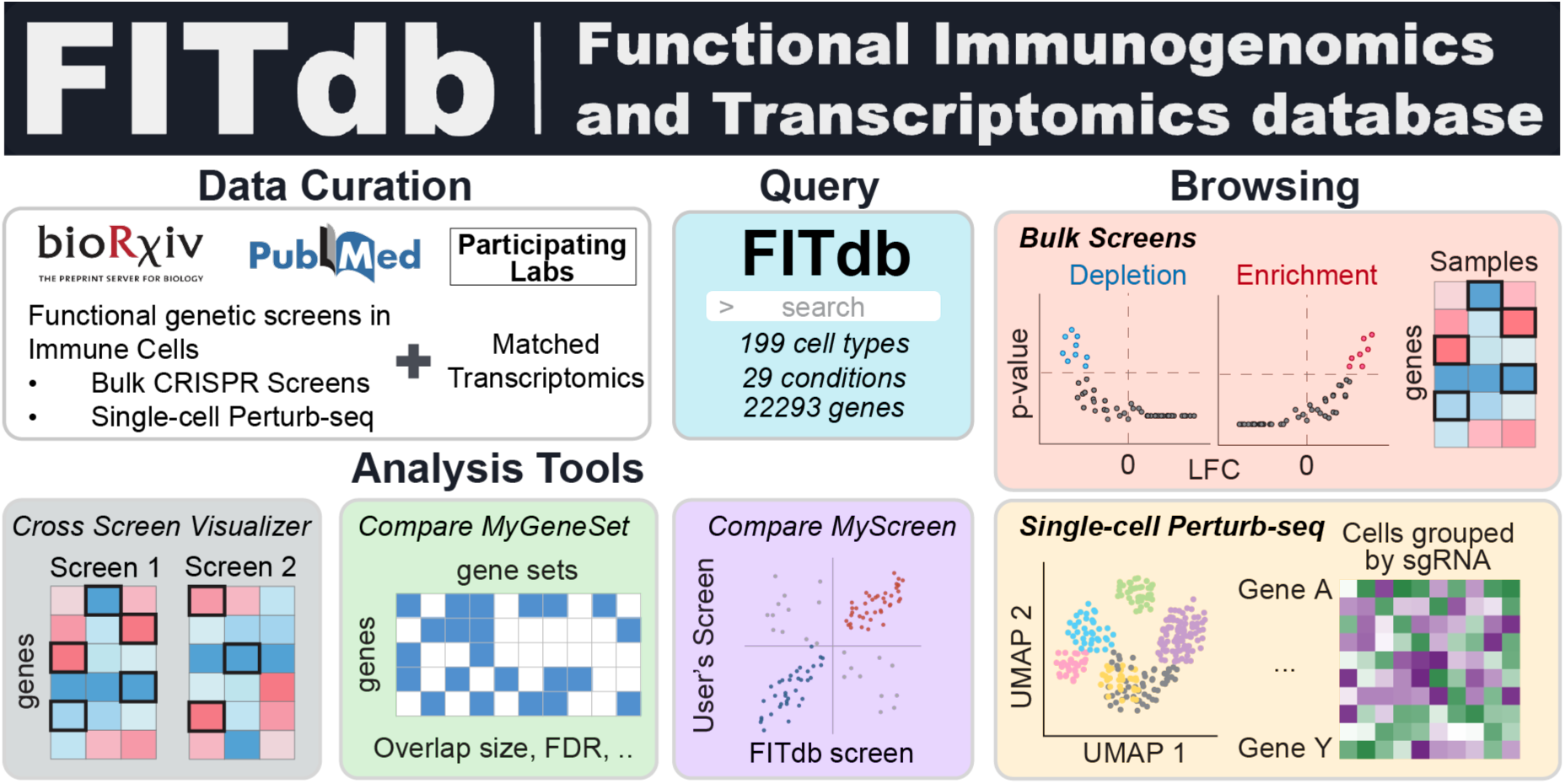

## Introduction

Current gene-editing technologies based on the CRISPR-Cas system provide a comprehensive toolbox for selectively and efficiently perturbing genes (1–3). Coupled with Next-Generation Sequencing (NGS), these tools enable the simultaneous interrogation of hundreds to thousands of genes in a single assay(1). The availability and cost-effectiveness of these approaches(4, 5) are driving a rise in functional genomics in immunology, providing an important causal mechanistic layer that complements transcriptional profiling(6–14). Hundreds of laboratories worldwide now leverage these tools to interrogate the vast, uncharted space of gene function across immune cell types and disease contexts. However, these studies are often targeted in scope, perturbing a pre-selected subset of genes, and are focused on specific cell types and conditions of interest. These experiments generate large, useful datasets, but these data remain largely unsearchable, limiting their accessibility, cross-study integration, and the systematic extraction of generalizable principles governing immune cell function. To bridge this gap, we have developed the **Functional Immunogenomics and Transcriptomics database** (**FITdb**), which curates and integrates functional genomics screens of mouse and human immune cells, coupled with their gene expression profiles, delivered through an intuitive, freely accessible web-based user interface and a suite of analysis and visualization tools.

Pooled functional screens rely on a relatively simple yet powerful metric, enrichment or deletion, typically shown as a Log_2_ Fold-Change (LFC) linked to a statistical value (p-value). To obtain these values, the frequencies of cells expressing a single-guide RNA (sgRNA) and a Cas9 protein are compared before (*input*) and after (*output*) the cell mixture is subjected to a selective pressure (*assay*)(5, 15) (**Figure 1A**). For instance, the initial composition of sgRNAs in an input population of activated CD8 T cells is compared to the sgRNA composition after these cells are adoptively transferred to a mouse bearing a tumor and recovered from the tumor-draining lymph node(16). Additionally, different outputs can be compared among them to assess the relative functional importance of each gene, for instance, in proliferating vs non-proliferating cells(17). For any given assay (tumor, infection, activation, etc), each gene either has a positive, negative, or non-significant regulatory role (**Figure 1B**). This role is interpreted based on the direction of the enrichment and the nature of the gene perturbation. That is, in the context of targeted gene deletion, such as with Cas9 (genetic cleavage) or iCas9 (transcriptional interference), significant enrichment of sgRNAs in outputs vs inputs (positive LFC) reveals negative regulators, whereas significant depletion of sgRNAs (negative LFC) reveals positive regulators (**Figure 1B**). In the context of CRISPR activation (CRISPRa) modalities, positive and negative regulators will show positive and negative LFCs, respectively(6). Thus, **positive regulators** are genes that promote the trait being selected for, while **negative regulators** are genes that restrict that trait for the given assay (**Figure 1B**). In summary, the sgRNA enrichment/depletion metric is universal, but the interpretation of the gene’s function depends on the assay design and the exact perturbation strategy. With this in mind, we have designed FITdb to provide an intuitive, biological interpretation of each screen.

**Figure 1.**
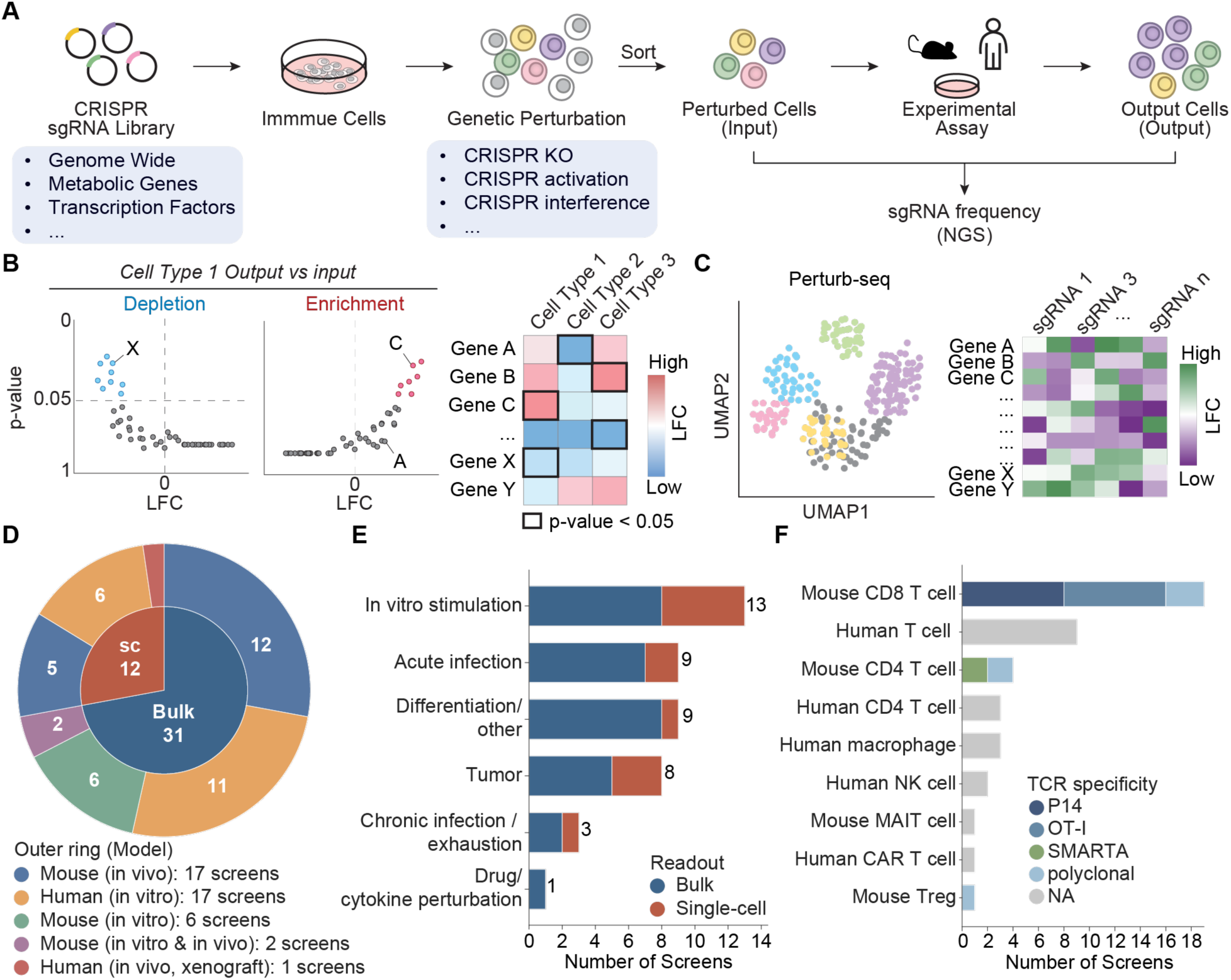
Overall design of FITdb. **(A)** Schematic of a typical pooled CRISPR screen included in FITdb. sgRNA libraries (genome-wide or focused) are introduced into immune cells to generate perturbations by CRISPR knockout, interference, activation, or others. Perturbed input cells are sorted, subjected to an experimental assay in mouse or human systems, and the output cells sequenced to quantify each perturbation’s effect. The sgRNA frequencies are measured in outputs and inputs for quantification of enrichment/depletion. **(B)** Readout of bulk pooled CRISPR screens, which measure perturbation effects on population sgRNA frequencies in output vs inputs, tested for depletion (left volcano plot) or enrichment (right volcano plot) in Cell Type 1 vs input. LFCs for each Cell Type can be combined into a heatmap plot where significant genes (adjusted p-value < 0.05 are highlighted with thick border line). **(C)** Readout of single-cell (Perturb-seq) screens, which measure perturbation effects on single-cell transcriptional states. Left: UMAP embedding of single-cell transcriptomes colored by sample. Right: schematic of per-perturbation LFC across genes, grouped by sgRNA identity. **(D)** Composition of the current FITdb collection by readout and experimental model. Inner ring: readout type, distinguishing bulk enrichment/depletion screens from single-cell transcriptional-state screens. Outer ring: experimental model system. Wedge size is proportional to the number of screens; counts are indicated. sc, single-cell **(E)** Number of screens per immunological challenge category, colored by readout type (bulk vs. single-cell). **(F)** Number of screens per cell type, colored by TCR specificity.

Single-cell CRISPR-based functional perturbations (perturb-seq) are enabled by simultaneous identification of the sgRNA and the cell’s transcriptome(18) (**Figure 1C**). These datasets often lack an input reference, although relative changes in sgRNA frequencies can be inferred by assuming equal starting populations or by comparison with non-targeting or scrambled sgRNA controls(1, 19). From these relative changes, we estimated and incorporated the effect of each perturbation on cellular fitness for each of the curated single-cell CRISPR screens in FITdb. Importantly, these assays enable the identification of perturbation-dependent gene expression changes, which are easily browsable in FITdb and via a third-party extension to the UCSC Cell Browser(20) (**Figure 1C**). FITdb contains single-cell functional screens using multiple modalities, including gene disruptions(21–26) and gene overexpression(27). These single-cell level screens typically target fewer genes than pooled screens, although recent studies report genome-scale screens in primary human T cells in vitro(28).

In summary, leveraging genome-scale bulk screens with high-resolution single-cell perturbation approaches is essential to capture both the breadth and mechanistic depth of the genetic programs that regulate immune cell function, and FITdb provides a unified resource for exploring both modalities.

### Database Content and Construction

A list of pooled and single-cell functional genetic screens of immune cells was compiled from a literature survey and dataset curation. **43** screens with available data were selected, of which **40** (from **29** independent peer-reviewed publications) have now been incorporated, in addition to **3** unpublished screens generated in our laboratory or by collaborators (**Supplementary Table 1**) (**Figure 1D**). Approximately one-fourth of the screens are in a perturb-seq format, including both overexpression and gene disruption (**Figure 1D**). Combined, these screens cover a diverse set of mouse and human immune cell types subjected to a range of immunologically relevant perturbations in *ex vivo*, *in vivo* and *in vitro* assays (**Figure 1E and 1F**). FASTQ were used as a starting point for re-analysis whenever possible. When these were unavailable, sgRNA count tables generated by MAGeCK count subcommand (v0.5.9.5) were used as a starting point(29). sgRNA- and gene-level frequency data for outputs vs inputs (*’Cell vs Input*’) and relevant output comparisons (*’Cell vs Cell’*) were used to build a relational database in MariaDB accessed via the Prisma ORM. Perturb-seq sgRNA assignments follow the author’s assignments. Dimensionality reductions, when not provided by the authors, are computed with the Scanpy (v1.11.1) workflow (pp.normalize_total(), pp.log1p(), tl.pca(), pp.neighbors, tl.umap())(30). The website was developed using SvelteKit and containerized with Docker, orchestrated via Docker Compose, with images built and deployed through a GitLab CI/CD pipeline. Interactive visualizations were implemented using Chart.js, extended with the chartjs-chart-matrix, chartjs-plugin-annotation, and chartjs-plugin-zoom plugins for heatmaps, pathway plots, and zoomable views. The interface incorporates client-side multi-select and typeahead filtering to enable rapid exploration of large datasets. Database queries are optimized through primary-key-indexed lookups scoped per screen/dataset, with a singleton Prisma client reused across requests to avoid connection overhead. Source code is maintained using Git, hosted on a self-managed GitLab instance. The website has been designed using UnoCSS, a Tailwind-compatible utility-first CSS framework, to provide a consistent, responsive experience across desktop, tablet, and mobile devices.

### Database Features and Web Interface

FITdb’s landing page enables a quick start in the main search bar with typeahead (**Figure 2A**). Entering a gene query directs the user to the “*Cross-Screen Visualizer*” page, where results for the queried gene are displayed across all screens available on the website (mouse by default). Underneath the search bar, four buttons provide direct links to the main tools of FITdb: “*Explore X Screens*” (Screen Browser), “*Cross-screen visualizer*”, “*Compare MyGeneset*”, and “*Compare MyScreen*” (**Figure 2A**). The bottom left provides a welcome message, contact information, and contributor acknowledgments, while the bottom right side includes a short video tutorial. Finally, a ReadMe button in the bottom left corner directs the user to a new page that provides an Introduction, Interface overview, Frequently Asked Questions (FAQ), and References.

**Figure 2.**
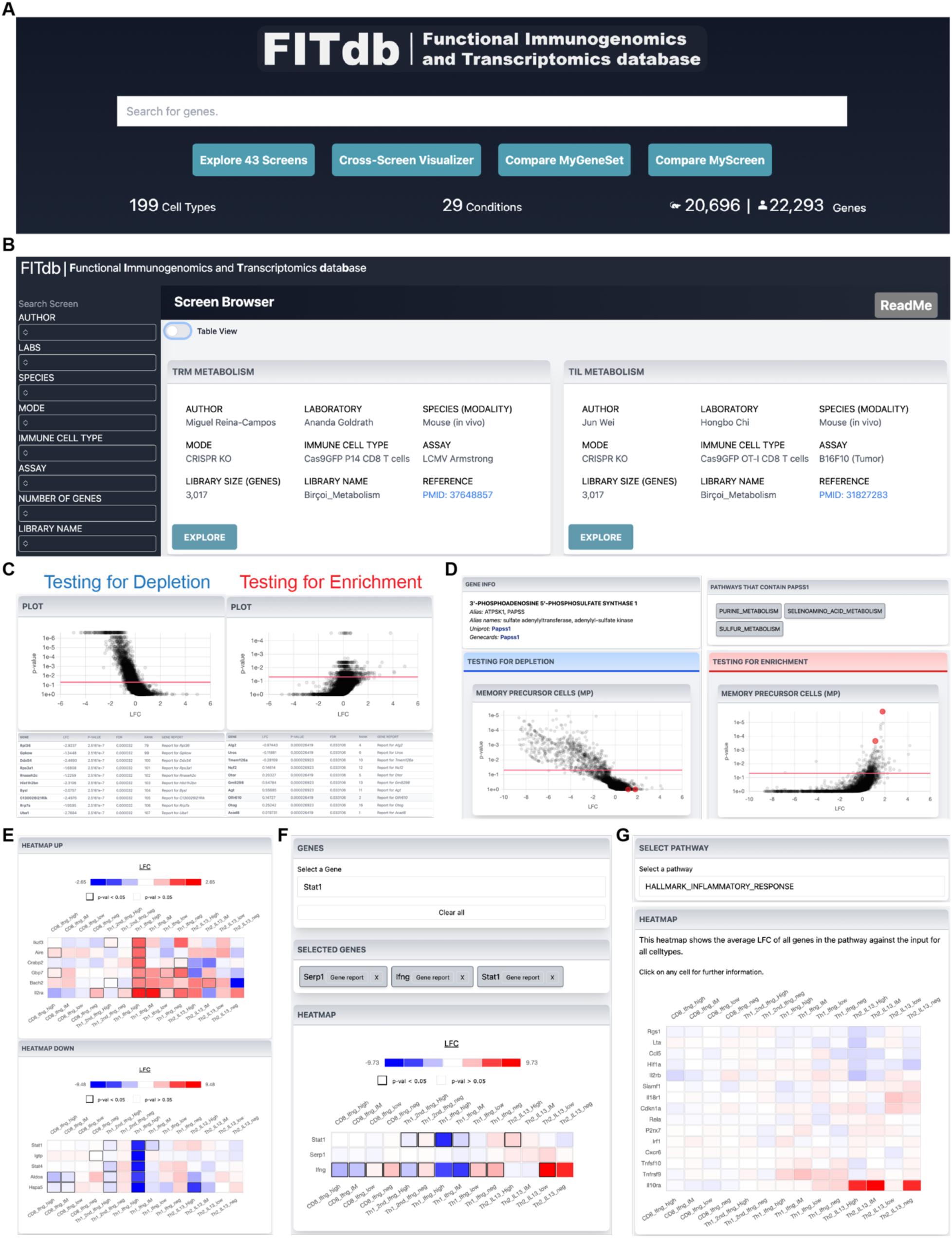
Analysis and visualization of individual bulk CRISPR screens. **(A)** Landing page of FITdb contains a gene search bar, and buttons leading to each of the specific functionalities: “*Explore X Screens*” (Screen Browser), “*Cross-Screen Visualizer*”, “*Compare MyGeneSet*”, and “*Compare MyScreen*”. **(B)** Overview of Screen Browser page. Left-sided menu allows to filer screens by authorship (first author and lab), species (human or mouse), mode (bulk vs single-cell, CRISPR KO/a/i), immune cell type, experimental perturbation, and size and name of the library. The right side contains screen metadata information about the screen, displayed either as a collection of cards or a table that can be switched with the toggle in the upper left corner. **(C)** Results display for a cell vs. input or cell vs. cell comparison, LFC and p-value for each gene in either direction (depletion and enrichment) is displayed as both an interactive scatter plot and a searchable table. **(D)** Gene report page. User can search/select a gene to view its detailed information, including pathway membership, aliases, and links to protein and gene databases. Its enrichment values (LFC and p-value) are also displayed as a scatter plot for each cell type or condition present in the study. **(E)** Heatmaps (split by enrichment or depletion) for all statistically significant genes across all cell types or conditions present in the study. Heatmap box is colored by LFC and statistically significant results are marked by a thicker black box. **(F)** The Custom heatmap page allows user to select genes of interest and visualize their LFC and p-values. **(G)** Under the Pathway analysis tab, the user selects a MSigDB hallmark gene set (e.g. HALLMARK_INFLAMMATORY_RESPONSE), and the resulting heatmap displays the LFC of each gene in that set, restricted to genes present in the screen’s library, across all cell types or conditions.

The Screen Browser, accessible via the “*Explore X Screens*” button under the search bar (**Figure 2B**), provides users with an intuitive entry point into FITdb. All screens are shown as a table by default (or card view if toggled, which is optimized for mobile), which can be filtered using the dark-themed side-bar panel using multiple metadata parameters, including Author, Laboratory, Species, perturbation mode (Mode), Immune Cell Type, Assay, Number of Genes, and library name to quickly locate specific datasets (**Figure 2B**). Screen metadata, including a direct link to the published reference via its PMID, is shown for each screen. Clicking on the “Explore” button on any screen of interest (card view) or on the Title link (table view) opens a new page for dataset exploration.

Each screen can be explored in detail through a dedicated interface that provides a suite of analysis modules tailored to its scope. These modules include: “*Cell vs. Input”*, “*Cell vs. Cell”*, “*Gene Report”*, “*Gene Heatmap”*, “*Custom Heatmap”*, and “*Pathway Analysis”* (**Figure 2C-G**).

The “*Cell vs. Input*” and “*Cell vs. Cell*” modules enable direct comparisons between immune cell populations to infer gene-specific regulatory roles. In “*Cell vs. Input”* analyses, the perturbed (*output*) population is compared with the initial *input* population to identify genes whose perturbation alters cellular abundance, as reflected in the depletion or enrichment of sgRNAs. In “*Cell vs. Cell”* analyses, two output populations are compared to uncover sample-specific functional differences. For each comparison, FITdb generates interactive visualizations that distinguish between depletion signals (sgRNAs reduced after selection) and enrichment signals (sgRNAs increased after selection), enabling rapid interpretation of gene function across conditions (**Figure 2C**). The “*Gene Report*” module provides an in-depth, gene-centric view integrating functional screening and transcriptomic data. Users can query individual genes to access sgRNA-level statistics, including LFCs, sequence information, and gene rankings across screen libraries. These data are complemented by gene expression profiles across immune cell subsets, facilitating interpretation within physiological contexts (transcriptomics is still only available in some bulk screens). Additionally, the platform incorporates curated annotations, such as Pathway Ontology gene sets containing the given gene, and direct links to external resources, such as UniProt(31) and GeneCards(32), to provide additional functional and pathway-level insights (**Figure 2D**). The “*Gene Heatmap*” and “*Custom Heatmap*” modules enable visualization of LFC enrichment/depletion values for each cell type (output vs input) across the cell types or conditions included in a given screen (**Figure 2E**). The *”Gene Heatmap*” automatically displays the values for all significant genes for a user-selected immune cell population, while the “*Custom Heatmap*” enables users to input predefined or custom gene lists and generate interactive plots. These heatmaps support dynamic exploration, for instance; clicking on any given heatmap cell in the plot will trigger a pop-up window prompting the user to explore that gene or cell type in detail in the “*Gene Report*” or “*Cell vs Input*” tabs, respectively. Additionally, plots can be exported as high-resolution “.svg” files for downstream use (**Figure 2E and 2F**). Finally, the “*Pathway Analysis*” module integrates CRISPR screening data with curated biological pathways to provide systems-level insights (**Figure 2G**). Users can interrogate specific pathways and visualize the distribution and behavior of constituent genes across screens (**Figure 2G**). Pathway gene sets were obtained from the Molecular Signatures Database (MSigDB) hallmark gene set collection(33) and from previous publications(9). Outputs include pathway-level summaries, gene-specific heatmaps, and classification of pathways enriched for negative or positive selection. Collectively, these features enable hypothesis generation about the molecular networks that govern immune cell function. (**Figure 2G**).

The “*Cross-Screen Visualizer*” enables integrated analysis of gene function across multiple CRISPR screens, immune cell types, and assays (**Figure 3A**). Users can input custom gene lists and select screens from the mouse or human screens to identify shared patterns or context-specific effects, facilitating target prioritization. Results are presented as an interactive heatmap of LFC values and statistical significance, with color-coding for depletion (negative LFC) and enrichment (positive LFC). This format enables rapid comparison of gene behavior across immune cell subsets and experimental conditions, helping distinguish consistent regulators from context-dependent effects (**Figure 3A**).

**Figure 3.**
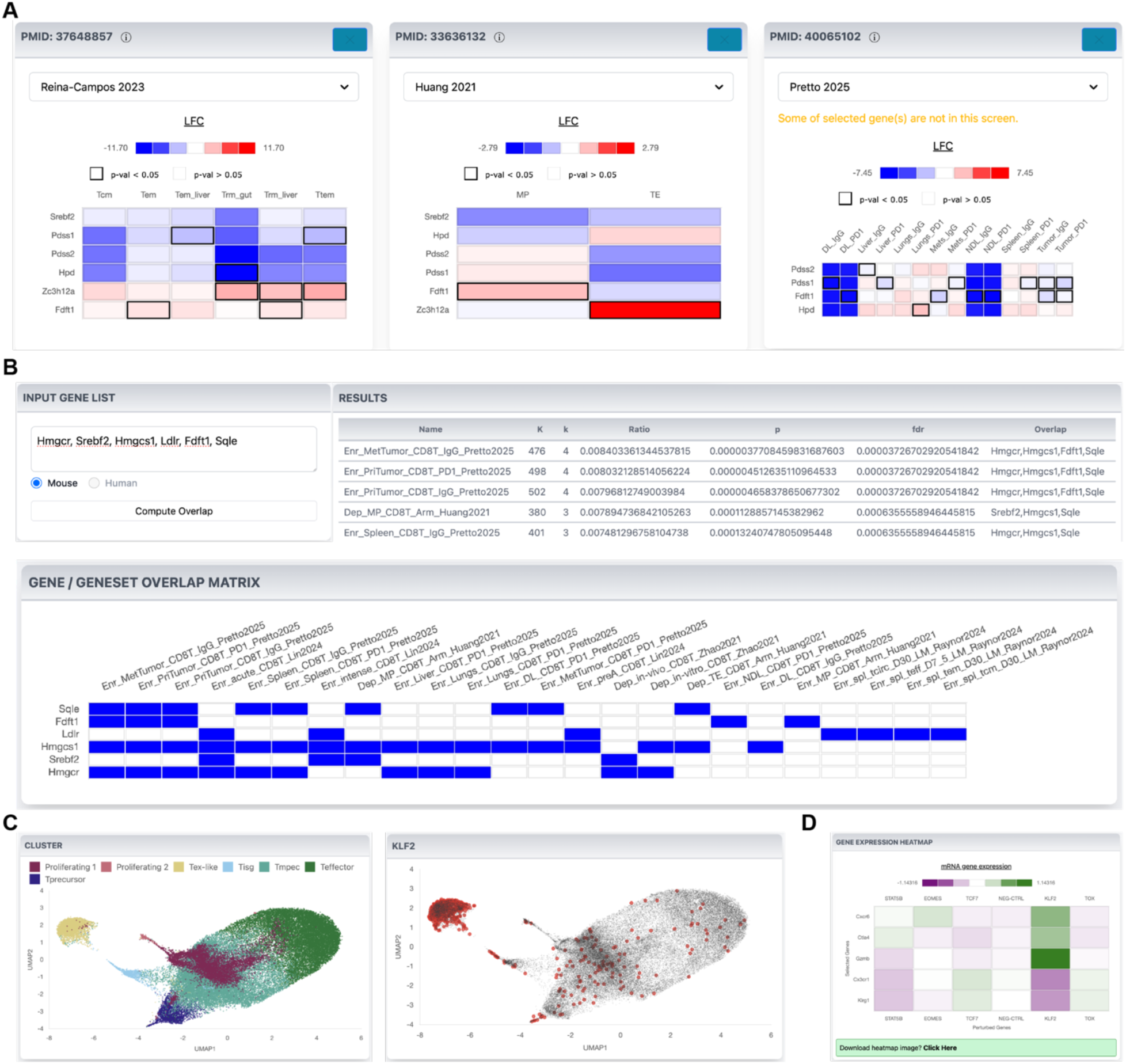
Analysis and visualization across bulk CRISPR screens and single-cell CRISPR datasets. **(A) “***Cross-Screen Visualizer*” generates LFC heatmaps for user-provided genes across all bulk screens in the collection containing at least one of the genes; screens missing a queried gene simply omit it rather than being excluded, and statistically significant results (p-val < 0.05) are marked by a box. **(B) “***Compare MyGeneSet*” tests a user-input gene set for over-representation against the curated FITdb gene sets using a hypergeometric test, returning overlap size, p-value, and False Discovery Rate (FDR). **(C)** Single-cell CRISPR dataset browser. Left: distribution of cells by cell type on the UMAP embedding. Right: distribution of cells by the gene being perturbed on the UMAP embedding (KLF2 sgRNA is shown). (**D**) Downloadable gene expression heatmap of selected genes (rows) in cells grouped by selected perturbed genes (sgRNA, columns).

The “*Compare MyGeneSet*” module allows users to find overlaps between a custom gene list and any of FITdb’s curated gene sets - the significantly enriched and depleted fitness genes identified per cell type across the curated CRISPR screens (**Figure 3B**). After entering a comma-separated gene list and selecting the relevant model, the tool computes the overlap between the input and each curated set and tests it for statistical over-representation, supporting hypothesis generation, validation of CRISPR screen hits, and contextualization within immunology-focused screens. Enrichment is assessed with a one-sided hypergeometric test (upper tail), where the sampling universe defaults to the union of all curated FITdb gene sets but can be restricted to a user-supplied background for the query, the database, or both; the overlap count, curated-set size, and this universe together determine each p-value. Results include every curated set sharing at least one gene with the input (this minimum overlap is configurable), each annotated with the overlapping genes, the overlap size, and a Benjamini-Hochberg False Discovery Rate (FDR), ranked so that the most significant sets appear first. An accompanying binary matrix reports membership between the curated gene sets and the input genes, enabling visualization of which input genes drive overlap across datasets (**Figure 3B**).

The “*Perturb-seq Browser”* provides an interactive interface to explore curated single-cell perturbation datasets in immune cells (**Figure 3C**). Users can visualize cellular embeddings, assess perturbation distributions, and examine transcriptional responses to specific genetic perturbations (**Figure 3C and 3D**). The platform also links each dataset to a UCSC Cell Browser session for deeper, single-cell– level exploration. The browser includes three main visualization tabs. The first is a scatterplot in which each point is a single cell, positioned by uniform manifold approximation and projection (UMAP), a dimensionality-reduction method that places transcriptionally similar cells close together in two dimensions(34). The UMAP view displays cell type or cluster annotations by default, with the option to overlay specific perturbations for direct comparison (**Figure 3C**). The bar graph summarizes the relative abundance of perturbed cells, shown as LFC relative to control conditions. The heatmap tab displays gene expression changes as Z-scores across perturbations, with options to customize gene and perturbation selection (**Figure 3D**). Together, these tools enable rapid interrogation of perturbation effects on immune cell states and gene expression programs at a single-cell level.

Finally, the “*Compare MyScreen”* module enables users to query their own bulk CRISPR screen results against curated datasets in FITdb. This tool identifies shared hits and unique findings, supporting validation and interpretation of experimental results by comparing user-provided data (as MAGeCK outputs) with reference screens. It also provides a statistical assessment of overlap strength and generates publication-ready visual outputs for direct download. The output includes interactive summaries of depletion and enrichment, with ranked tables reporting LFC, raw p-values, and FDR. A gene overlap module quantifies shared hits between datasets and provides a scatter plot comparing LFC values across screens, with adjustable FDR thresholds for filtering. In addition, side-by-side comparison tables summarize enrichment and depletion statistics for overlapping genes.

### Use case example: Functional role of downstream TCR components

T cell receptor (TCR) engagement initiates an intracellular signaling cascade that connects antigen recognition at the cell surface to transcriptional programs driving cytokine production and proliferation(35). A central component of this process is the transmembrane adaptor LAT, which recruits signaling proteins downstream of the TCR and promotes activation of the Ras-MAPK pathway(36). LAT2 (also known as LAB) is a related but less well-understood adaptor implicated in the regulation of lymphoid and innate immune signaling(37). We used FITdb to interrogate the functional effects of LAT and LAT2 across complementary CRISPR datasets. Using the “*Cross-Screen Visualizer*”, we queried *LAT* and *LAT2* across human immune-cell bulk screens (**Figure 4A**). In the ‘Schmidt 2022’ CRISPR activation and interference screens in T cells, these genes showed significant opposing effects in both modalities (**Figure 4A**). That is, in the activation screen, LAT promoted enrichment of CD4 T cells with high IL-2 production and CD8 T cells with high IFN-γ production, consistent with its established role as a positive regulator of TCR signaling. In contrast, LAT2 activation depleted these cytokine-high populations, suggesting that LAT2 restrains productive T cell activation despite its structural and functional relationship to LAT. Consistent with these roles, the interference screen showed LAT-targeting cells were significantly depleted from the CD4 IL-2 high and CD8 IFN-γ high populations, while LAT2-targeting cells were significantly increased in the CD8 IFN-γ high population (**Figure 4A**). Querying mouse bulk screens showed *Lat2* to have a negative suppressor role in T cell fitness under various conditions (‘Lin et al 2024’) (38) (**Figure 4B**). These data can also be explored under the “*Gene Report*” page for the respective datasets, which also provides the underlying sgRNA sequences, effect sizes, and statistical values. Next, we examined the Schmidt 2022 Restimulation Perturb-seq dataset to investigate how the activation of either adaptor impacts T cell differentiation, proliferation, and the expression of cytotoxic molecules. Looking at the distribution of LAT-vs LAT-2 sgRNA-containing cells in the UMAP embeddings showed an enrichment of LAT2 cells in the Negative Regulators Group (**Figure 4C**). Plotting gene expression in cells grouped by LAT and LAT2 activation revealed a differential effect on the expression of several granzymes (*GZMA*, *GZMB*, and *GZMK*) and cell proliferation genes, consistent with the putative antagonistic functions of these genes in T cell fitness (**Figure 4D**). These distinct responses suggest compensatory upregulation of alternative signaling components following disruption of either adaptor, which can be explored further following the link to the UCSC Cell Browser. Finally, to understand potential upstream transcriptional regulators of *LAT* and *LAT2* we utilized a second perturb-seq examining gene circuits controlling T cell rest and activation(24) (**Figure 4E**). Plotting gene expression of *LAT* and *LAT2* across the 28 perturbations included in this perturb-seq showed overall distinct putative regulators, including *MYC* and *DNMT1* as unique inhibitors of *LAT* and *LAT2*, respectively, and *MED12* and *SOCS3* as promoters of *LAT*. Together, this example illustrates how FITdb can connect cross-screen functional phenotypes with sgRNA-level evidence and perturbation-dependent transcriptional responses to generate mechanistic hypotheses.

**Figure 4.**
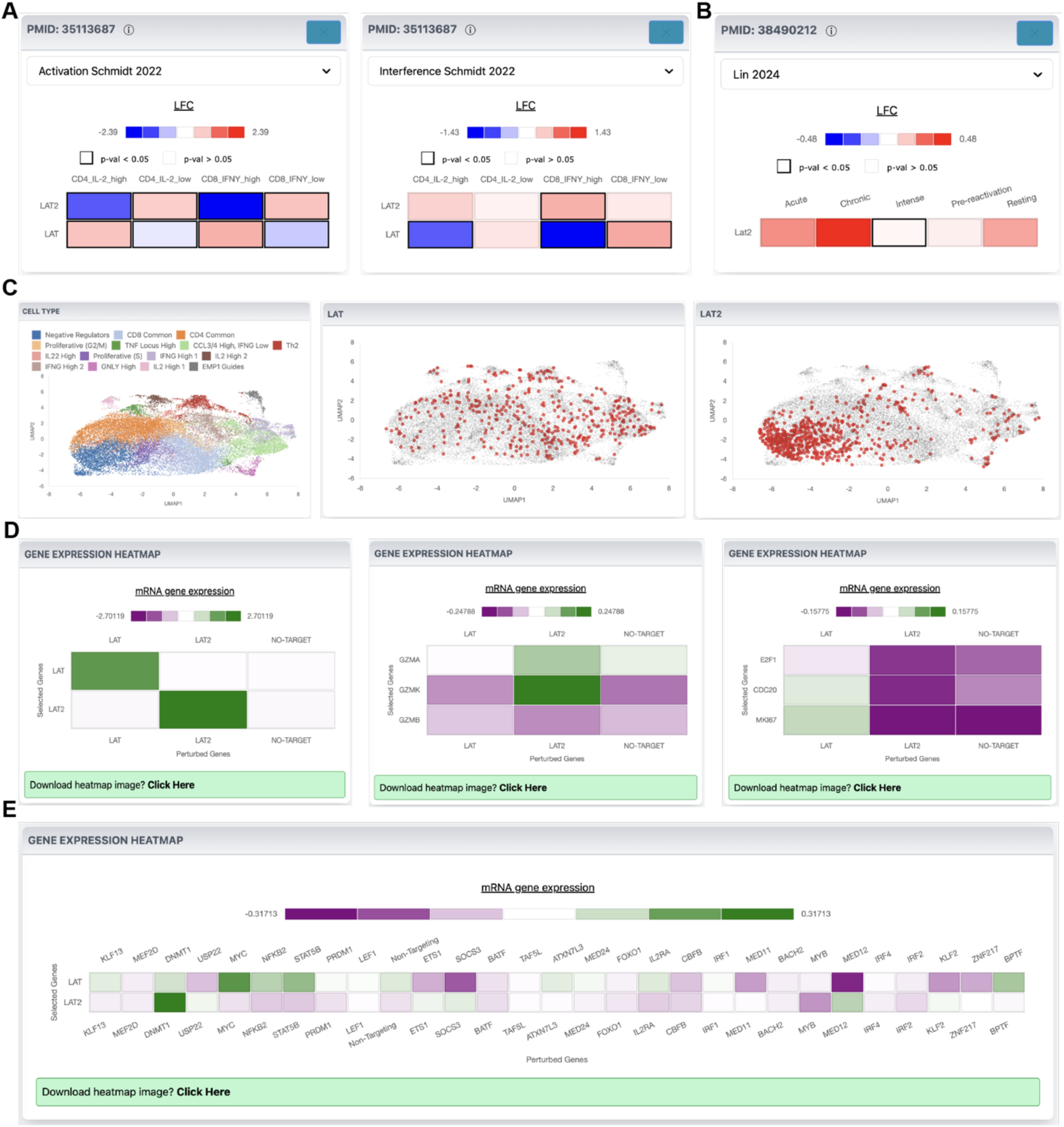
Interrogation of LAT and LAT2 functions in T cells. (**A**) ‘*Cross-screen comparison*’ results for LAT and LAT2 queries in human and (**B**) mouse T cell datasets. (**C**) UMAP embedding of the CRISPR activation Perturb-seq dataset ‘Schmidt 2022 Restimulation’ including LAT and LAT2 perturbations in T cells colored by cluster name (left), LAT sgRNA (middle), and LAT2 sgRNA (right). (**D**) Expression of indicated genes (rows) for cells grouped by their respective perturbation (LAT and LAT2 (columns) in ‘Schmidt 2022 Restimulation’ Perturb-seq. (**E**) Expression of LAT and LAT2 in human primary CD4 T cells grouped by perturbation in the ‘Arce 2025’ Perturb-seq.

### Comparison to Existing Resources

Existing functional genomics databases include comprehensive resources, such as BioGRID ORCS (https://orcs.thebiogrid.org/), and targeted resources for specific cell types, such as cancer cells (DepMap, https://depmap.org/portal/), or specific contexts, such as genes related to lipid biology (CRISPRlipid, https://crisprlipid.org/). Two other databases cover functional genome-wide immunogenomics of T cells in the oncology setting: TCPGdb (http://tcpgdb.sidichenlab.org/) and iCRAFT(39) (https://icraft.pku-genomics.org/#/homepage). A third more focused database exists for metabolic regulators of T cells (40, 41) (https://functionalimmunogenomics.shinyapps.io/crispr/). Our database resource is positioned to cover immunogenomics and transcriptomics of immune cells for any CRISPR modality, including pooled and single-cell, of any size, in mouse and human across any disease or perturbation setting. To our knowledge, FITdb is the only resource to integrate both bulk and single-cell perturb-seq datasets with a suite of tools for browsing, analyzing and comparing across experiments.

### Availability and Maintenance

FITdb is actively maintained, with new publicly available CRISPR screening datasets curated and incorporated on a regular basis. The application is deployed using Docker on dedicated internal LJI infrastructure, enabling streamlined updates and maintenance through a GitLab CI/CD pipeline that builds a container image on merge and deploys it via automated SSH rollout to the target host. Since the public release of the beta version in January of 2025, we have pushed 7 major updates, approximately once every three months, adding new screens and functionalities. Based on our growth projections, we anticipate adding ∼10-30 screens per year. Contact information is provided on the landing page for reporting issues, suggesting new features, or submitting datasets for incorporation into FITdb.

### Conclusion and Future Plans

We have created FITdb to provide a freely accessible, integrated compilation of in vitro, in vivo, and ex vivo functional screens of mouse and human immune cells. In its latest update (as of July 20^th^ of 2026) it includes 43 independent screens, with 199 immune cell types, states, or treatment conditions, covering 20, 696 mouse and 22, 293 human genes and 5 different CRISPR modalities (**Figure 1D-F**). We currently have 31 curatable screens pending database incorporation and, based on previous publication records, expect new screens at a rate of 2 per month. At the current rate, we expect FITdb will host more than 80 screens within the end of 2027

We envision several additional tools and enhancements will be added to FITdb during 2026-2027. First, we envision developing a gene sgRNA picker tool informed combining sequence prediction and actual experimental data. We also plan to use the compiled FITdb dataset to develop a statistical imputation method to predict gene function for unseen genes (not included in the screening library). Furthermore, we plan to add transcriptomic data covering most immune cell types and states via an Application Programming Interface (API) call to the Immunological Genome Project (ImmGen). Finally, we plan to provide our own API to enable users to run more complex, custom queries. Collectively, FITdb will provide a comprehensive and continuously evolving resource to accelerate discoveries in immunology.

## Data Availability Statement

All the re-reprocessed data shown in FITdb are available upon request.

## Acknowledgments

We thank the following principal investigators and their research groups for generating and sharing CRISPR screening datasets that contributed to the development of this database: Alexander Marson, Julia Carnevale, Hongbo Chi, E. John Wherry, Jeffrey C. Rathmell, Marcela V. Maus, Arlene H. Sharpe, Crystal L. Mackall, Neville E. Sanjana, Shimon Sakaguchi, Katayoun Rezvani, Ansuman T. Satpathy, Junwei Shi, Nikhil S. Joshi, Min Peng, Daniel S. Peeper, Mario Di Matteo, Massimiliano Mazzone, Wei Wang, Susan Kaech Aditya Murthy, and Deyu Fang.

